# Learning Epithelial Elasticity via Local Tension Remodeling

**DOI:** 10.64898/2025.12.17.694921

**Authors:** Sadjad Arzash, Shiladitya Banerjee

**Affiliations:** School of Physics, Georgia Institute of Technology, Atlanta, GA 30332

## Abstract

Biological materials, like epithelial tissues, exhibit remarkable adaptability to mechanical stresses, dynamically remodeling their structure in response to external and internal forces. A key challenge is understanding how these tissues store a memory of past mechanical stimuli. Here, we investigate this memory using an active Vertex Model of epithelial sheets incorporating a local, mechanosensitive tension-remodeling rule where junctional tension updates depend on strain, acting as a slow, history-dependent variable. We demonstrate three hallmark mechanical consequences of this memory mechanism. First, a localized, short contractile cue permanently reprograms the global shear modulus, with the direction of change (stiffening or softening) controlled by the tension remodeling rate. Second, the tissue stores a long-range mechanical memory: a prior stimulus at one site modulates the tissue’s response to a subsequent, distant stimulus, mediated by coupling across the entire junctional network. Finally, we show that simple cyclic bulk deformation acts as a training protocol that autonomously tunes the tissue’s constitutive properties, including programming the Poisson ratio to auxetic (negative) values. These findings position epithelial mechanics within the framework of unsupervised physical learning, identifying the mechanosensitive remodeling rates as powerful control parameters for designing programmable tissue-scale rheology.

## INTRODUCTION

Many disordered materials achieve functional properties by storing memories of past stimuli [1–8]. Under repeated driving, these systems undergo irreversible microstructural changes so that their subsequent response encodes information about the training history [9–11]. A particularly relevant example is the “directed aging” protocol, in which disordered elastic networks are slowly driven while they relax, allowing local bond stiffnesses or rest lengths to evolve [2]. This algorithm can steer a network toward target mechanical tasks, such as producing an auxetic response, where the Poisson ratio is negative and the material expands laterally when stretched [12].

These ideas have recently been unified under a broader framework of “physical learning” in driven soft materials [13– 19]. In this view, a physical system is described by fast dynamical degrees of freedom, such as node positions in an elastic network, coupled to slow internal variables, such as bond stiffnesses or rest lengths, that serve as tunable/learning parameters. Learning consists of updating these internal variables according to local rules, with the goal to optimize a task-dependent cost function [20] Because each update uses only locally accessible information, the training is decentralized and does not require computing global gradients, in contrast to standard gradient-descent optimization in parameter space.

Epithelial tissues are a particularly important case study for this paradigm, representing biological materials that inherently possess adaptive, learning capabilities. As active, confluent sheets, their junctional tension, adhesion, and cortical forces are dynamically remodeled by both external and internal cues [21–29]. While passive mechanical models of epithelial tissues (which rely on fixed parameters) can reproduce baseline viscoelastic spectra and geometry-driven rigidity transitions [30–32], they are unable to capture history-dependent changes in stiffness, spatial patterning in force transmission, or mechanosensitive remodeling observed *in vivo* [33–35]. These biological processes demand material adaptation, making mechanosensitive remodeling rules essential for epithelial mechanics.

Adaptive models bridge this gap, explicitly placing epithelia within the physical learning framework. For instance, allowing cell shapes to tune as additional degrees of freedom shifts and controls rigidity transitions, while local, length- and orientation-dependent updates of edge tensions reproduce adaptive tissue flows during convergent extension[36, 37].

Critically, unlike supervised learning protocols, biological systems, such as epithelial tissues, do not employ an external cost function or protocol for adaptation. Their remodeling rules are driven intrinsically by local biophysical feedback mechanisms [34]. This places epithelial tissues in the category of unsupervised physical learning, where the system’s material properties are autonomously programmed by its inherent adaptive rule, rather than by a pre-designed task or global minimization scheme. We hypothesize that the mechanosensitive tension-remodeling rule, inherent to epithelial junctions, is sufficient for implementing unsupervised physical learning in tissue monolayers, enabling the encoding of long-range mechanical memory and the exhibition of autonomous programmability of global constitutive properties.

Building on our previously introduced framework of mechanical remodeling in epithelial junctions [28, 33, 34, 38, 39], we consider a model in which fast mechanical variables, such as cell vertex positions and cell pressures, relax quickly, while slower junctional tensions record memory through local, strain-dependent rules. Specifically, each junction’s tension is updated selectively in response to strain thresholds, with distinct rates for contraction (*k*_*C*_) and extension (*k*_*E*_). The ratio of these rates, *k*_*C*_*/k*_*E*_, serves as the key control parameter for the tissue’s mechanical programming.

We find that a short, localized contractile cue can rewire the tissue’s macroscopic mechanical properties; the shear modulus increases or decreases depending on *k*_*C*_*/k*_*E*_, highlighting this ratio as a sensitive control parameter for tissue-scale stiffening or softening. Furthermore, after an initial signal and full mechanical relaxation, subsequent stimuli at distant sites elicit responses distinct from näive tissues, demonstrating a history-dependent, long-range memory stored in the junctional network. Finally, subjected to small-amplitude, cyclic bulk driving, the tissue’s mechanical response becomes trainable, with the Poisson’s ratio tunable to auxetic (negative) values when *k*_*C*_ *> k*_*E*_. Conceptually, these findings position epithelial mechanics alongside other trainable materials: slow, local parameter adaptations, directed by rapid, globally driven fields, yield targeted changes without centralized optimization.

## METHODS

### Epithelial vertex model with mechanosensitive tension remodeling

We model epithelial tissue dynamics using a modified version of the standard two-dimensional vertex model [30, 40– 42]. In this framework, each cell is represented as a polygon, and the degrees of freedom are the positions of the vertices shared between neighboring cells. The mechanical energy of the tissue is given by

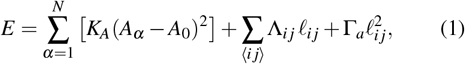

where the first summation runs over all cells, and the second over all edges between adjacent cells. Here, *A*_*α*_ and *A*_0_ are the current area and the target area of cell *α*, and the coefficient *K*_*A*_ control the stiffness associated with area deformations. The second term describes the energy due to interfacial tension: each cell-cell junction *i j* of length ℓ_*ij*_ is associated with an active tension Λ_*ij*_(*t*), which evolves dynamically. The last term describes the energy due to actomyosin contractility, where Γ_*a*_ is the contractile force per unit length of the edge [33, 38]. The total tension on edge *i j* thus becomes *T*_*ij*_ = Λ_*ij*_ + 2Γ_*a*_ℓ_*ij*_. This equation implies that the tension along a junction, arising from actomyosin contractility, is proportional to its length. This models the postiive feedback effect that myosin recruitment increases as junction length increases [43]. Position **r**_*i*_ of vertex *i* evolves in time as: 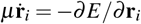, where *µ* is the vertex friction coefficient. Several recent studies have shown that intercellular tension in epithelial junctions is not fixed but continuously regulated by mechanochemical feedback [24, 28, 33–35]. To capture this adaptive behavior, we use the tension–remodeling model recently developed by one of us [33, 38], which quantitatively reproduces experimental observations of optogenetically driven junction contractions [33, 34]. In this framework, the tension Λ_*ij*_ along each cell–cell junction evolves in time according to its strain,

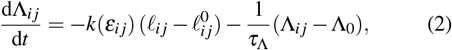

where ℓ_*ij*_ is the instantaneous junction length,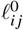 is its rest length, and the strain is 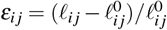. The second term describes tension relaxation to its equilibrium value Λ_0_ over a timescale *τ*_Λ_. The tension remodeling rate *k*(*ε*_*ij*_) depends on the strain as (see Figure 1a)

**FIG. 1.**
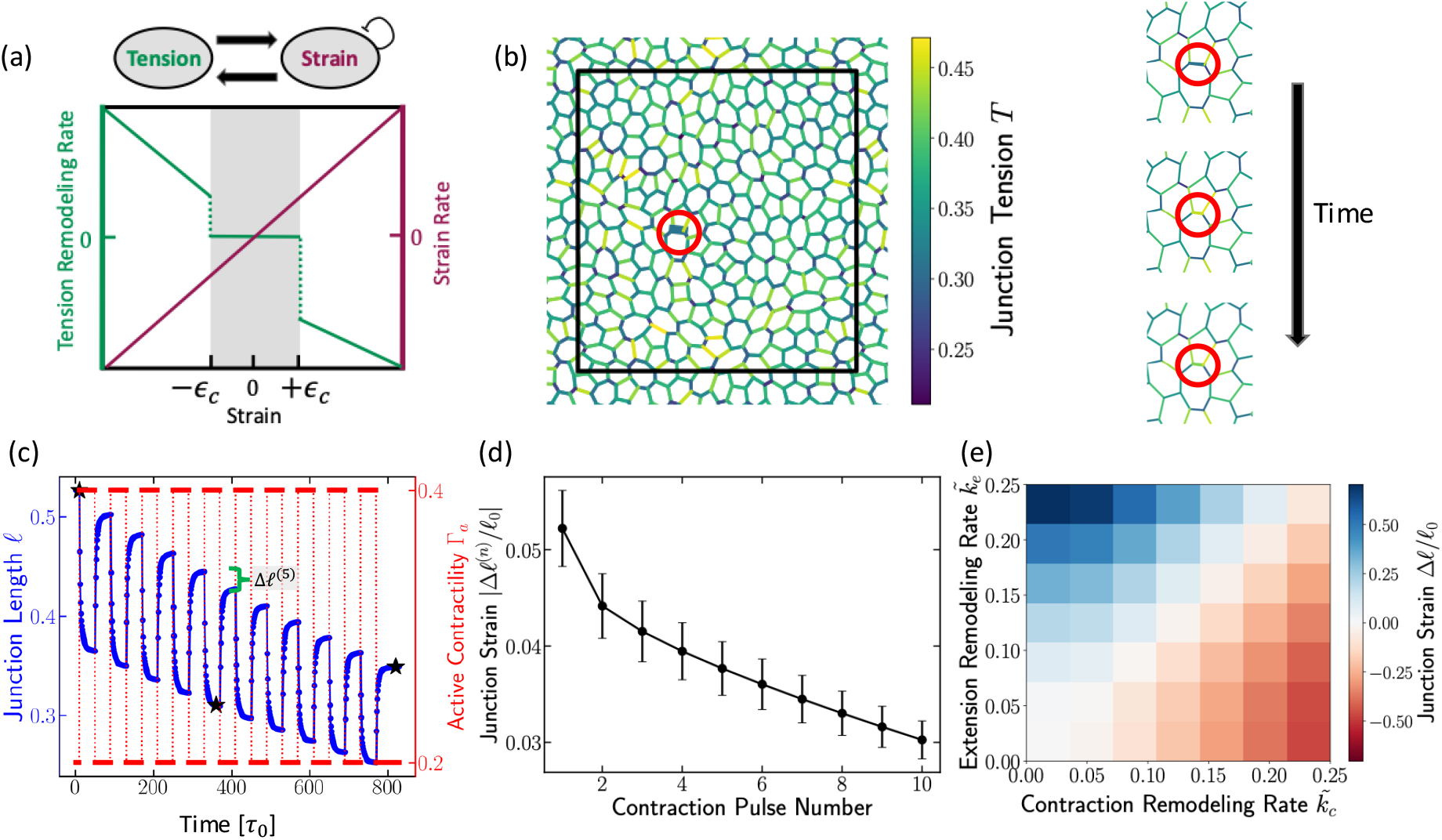
Adaptive dynamics of single junctions. (a) Schematic of the tension-remodeling model. In the contracted regime (ε < ε_c_), edge tension increases at a rate k_C_ while in the extension regime (ε > − ε_c_), the tension decreases at a rate k_E_. The edge strain relaxes continuously at a rate k_L_. (b) A randomly selected edge in the tissue, with the colorbar indicating edge tensions. On the right: Three zoomed-in snapshots highlight the same edge at different times during application of a contractile active signal; the corresponding time points are marked with stars in panel (c). (c) Time series of the selected edge length (blue) under pulsatile contractile activations (red), illustrating how the edge progressively shortens with each activation. (d) Net reduction in edge length as a function of the number of activations. Although repeated activations further reduce the edge length, the incremental reduction diminishes with each sequential activation, akin to habituation. (e) Phase diagram of junctional strain as a function of 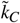 and 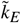 after the cessation of the contractile signal shown in red in panel (c).

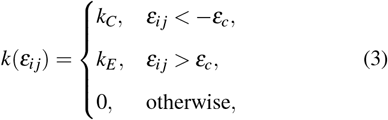

where *ε*_*c*_ is a local threshold strain for junction remodeling. For positive *k*_*C*_ and *k*_*E*_, tension increases under contraction at rate *k*_*C*_ and decreases under extension at rate *k*_*E*_. The threshold *ε*_*c*_ reflects the experimental observation that junctions remodel only above a critical strain amplitude, corresponding to sufficiently strong or sustained force [28, 33].

In addition to tension remodeling, cellular junctions continuously relax strain through rest-length remodeling. The rest length 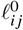 of each junction evolves toward its current length ℓ_*ij*_ at a rate *k*_*L*_ (see Figure 1a),

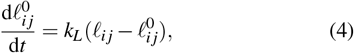

representing the gradual turnover of strained actomyosin structures and their replacement by unstrained ones. This relaxation process naturally arises from cytoskeletal turnover and myosin exchange within junctional networks [33].

An important consequence of Eq. 4 is that the tissue gradually forgets prior deformations over the characteristic timescale 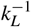. Sustained contractions therefore remodel junctions only up to a finite limit, as strain is progressively relaxed, whereas pulsatile contractions interspersed with rest periods enable cumulative, irreversible shortening through ratcheting [28, 33]. This interplay between tension remodeling and strain relaxation determines how long-term mechanical memory is written and erased in the tissue.

We perform our simulations using the open-source `cellGPU` package [44]. In the simulations, we nondimensionalize the equations of motion by setting *K*_*A*_ =1, *A*_0_ = 1, and *µ* = 0.1. Lengths are measured in units of 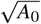, forces in units of 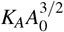, and time in units of the characteristic timescale *τ*_0_ = *µ/*(*K*_*A*_*A*_0_) With our parameter choices, one simulation time unit corresponds to a physical timescale of order ∼ 14 s [38]. To initialize the tissue, we place *N* random seed points in a square periodic box of size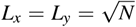 and construct the corresponding planar tissue using a Voronoi tessellation. To generate the initial tissue configuration, we first construct a confluent tiling governed by the energy *E* =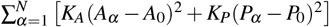, where *A*_*α*_ and *P*_*α*_ are the area and perimeter of cell *α*, and *A*_0_ and *P*_0_ are their target values [30, 31]. We set *K*_*A*_ = *K*_*P*_ = 1, *A*_0_ = 1, and *P*_0_ = 3.7, and obtain a mechanically equilibrated state by minimizing *E* with respect to vertex positions. The resulting force-balanced configuration serves as the reference tissue. We then initialize the edge tensions Λ_*ij*_ in our adaptive vertex model to match the tensions computed from this equilibrium configuration. We then run the overdamped dynamics of (1) together with tension dynamics (2) and rest-length dynamics (4) until the system reaches a steady state with uniform active tensions, Λ_*ij*_ = Λ_0_. This state serves as the initial tissue configuration for all subsequent simulations.

During all simulations, T1 transitions are triggered when an edge length falls below a prescribed cutoff value (see SI for details). In the results presented below, we work in the limit of slow tension relaxation compared to tension remodeling, 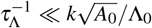. The effect of faster relaxation timescales *τ*_Λ_ is explored in detail in the Supporting Information, and its implications for the robustness of mechanical memory are discussed in the main text.

## RESULTS

### Single junction mechanics and memory encoding

We first elucidate the mechanism of mechanical memory encoding at the scale of a single cell-cell junction, which is crucial for the tissue’s emergent collective behavior. To this end, we examine the mechanical response of a single junction subjected to a pulsatile Γ_*a*_(*t*) (Fig. 1 b,c), simulating pulsatile contractions that are widely observed during morphogenesis [45– In the tension–remodeling model, these periodic stimuli produce a clear ratcheting effect: each pulse shortens the junction and the post-pulse length remains below its pre-pulse value, consistent with epithelial junction experiments [34]. Mechanistically, when the instantaneous strain crosses the negative threshold in (3), the update rule in (2) increases the junctional tension during the contracted phase. After the external contractility is removed, the junction relaxes toward a longer length, but the elevated tension (the stored memory) prevents full recovery, yielding a net shortening (Fig. 1c).

Successive pulses further increase tension and accumulate shortening, while the incremental shrinkage per pulse decreases over time (Fig. 1d), reflecting adaptation of the junctional state. This diminishing response under repetitive stimuli is a signature of *habituation*, a simple form of nonassociative learning [50]. In this context, habituation demonstrates that the junction possesses memory: the stored tension from previous pulses modifies the system’s current mechanical response, leading to a permanent, adapted state. This process validates the local tension-remodeling rule as a mechanism for encoding and accumulating mechanical memory.

The direction and magnitude of the permanent length change (Δℓ*/*ℓ_0_) are controlled by the two dimensionless tension remodeling rates, 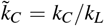 (contraction remodelling rate) and 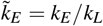 (extension remodeling rate). This ratio acts as the fundamental control parameter for local mechanical adaptation (Fig. 1e). When the contraction remodeling rate dominates 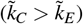, pulses drive a net shortening. Whereas, when the extension remodeling rate dominates 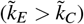, the same contractile pulses can surprisingly produce a net lengthening of the junction. In the SI, we show how an edge successively lengthens under a pulsatile contractile signal in the regime 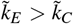. This behavior is summarized in the phase diagram of the junctional strain Δℓ*/*ℓ_0_ (Fig. 1e), which delineates the shortening and lengthening regimes in the 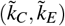 plane. The ratio 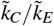 determines the direction of the final structural change. Moreover, the net edge length change depends sensitively on the pulse period: for short periods the length change is negligible, whereas increasing the period produces larger changes that eventually saturate at a plateau value for this signal (see Fig. S2 in the SI). Based on experimental data [33, 51], we expect epithelial tissues typically operate in the regime where the contraction-induced remodeling rate is higher 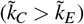, favoring net shortening and stiffening. The single-junction ratcheting and habituation establish the microscale learning rule that enables the emergent, collective mechanical reprogramming studied in the subsequent sections.

### Local cues reprogram global elastic property

We next ask whether localized, pulsatile contractility can reprogram tissue-scale material properties, demonstrating the macro-scale consequence of the junctional mechanical memory established previously. We apply contractile pulses to a circular region of radius *R* (Fig. 2a) and track how local junctional remodeling alters the tissue’s global mechanical response (Movie S1).

**FIG. 2.**
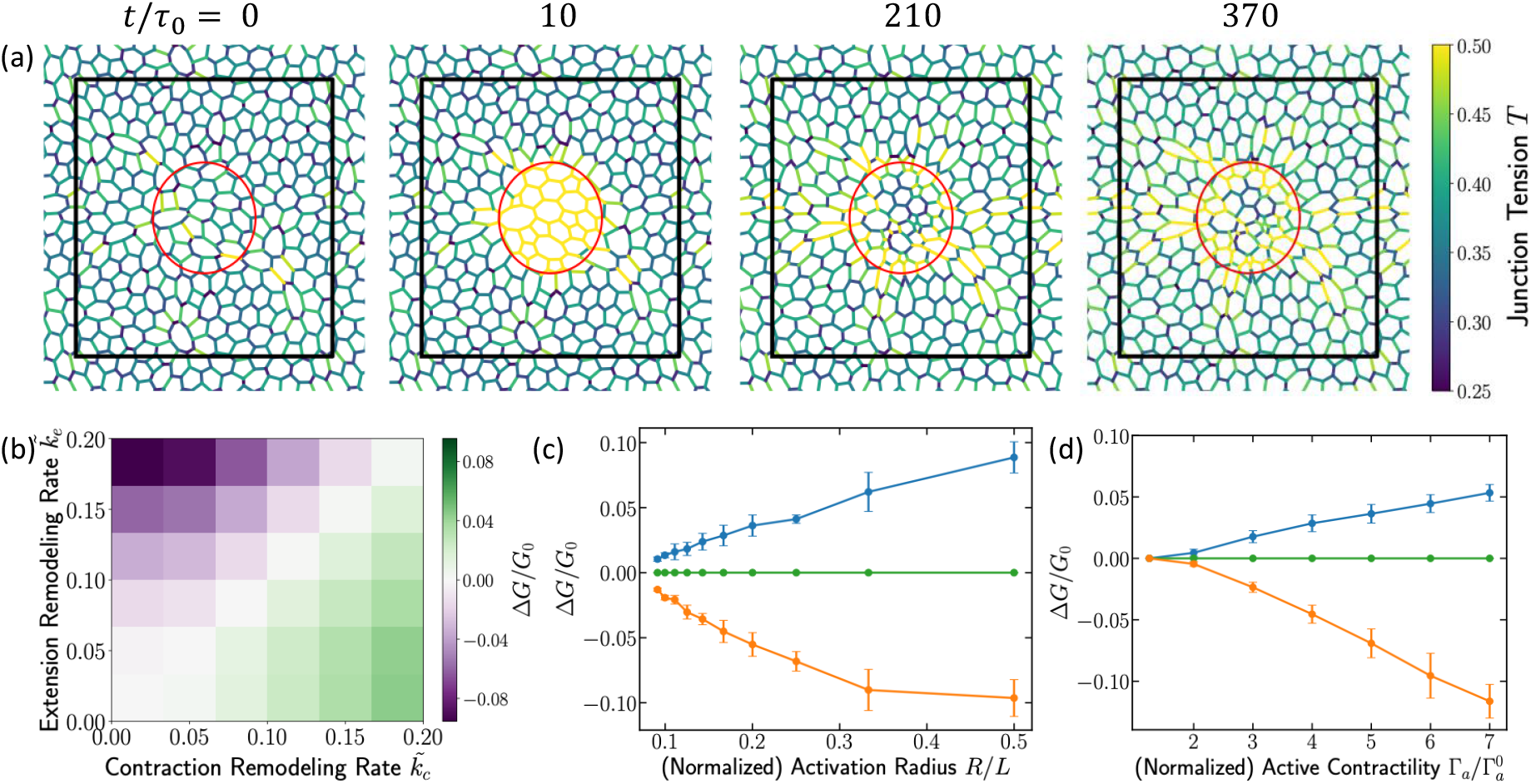
Local junction remodeling reprograms global mechanical properties. (a) Simulation snapshots of a circular activation region of radius R at the tissue center. The colorbar indicates edge tensions. The first snapshot shows the initial configuration with no contractile signal. The second snapshot shows the response during activation, where edges within the region develop high tension. The remaining snapshots illustrate the relaxation after the signal is removed, revealing that high-tension edges persist and spread into surrounding cells. These dynamics demonstrate that a locally applied contractile signal propagates through the tissue via feedback between tension and strain fields. (b) Phase diagram of the change in tissue shear modulus after five consecutive contractile activations applied to a central region of size R/L = 1/6. The applied signal matches that of Fig. 1, with 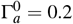 (low contractility), 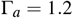 (high contractility), and pulse duration 40. (c) Change in shear modulus at 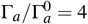. as a function of region size R/L. Blue: 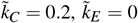; green: 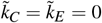; orange: 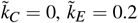 Change in shear modulus at R/L = 1/6 as a function of the contractility ratio 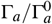. Color scheme as in (c).

Localized contraction pulses initiate a sequence of events that generates non-local changes in tissue properties via memory propagation. First, junctions within the activated region contract, and through the process of tension remodeling and strain relaxation, they increase their active tension and permanently reduce their rest length. These local changes encode a mechanical memory of the stimulus in the junctional network. This local contraction then induces strain in the neighboring cell junctions. Because tension remodeling is mechanosensitive, these strained neighboring junctions undergo tension remodeling, which, in turn, alters their tension and rest length, propagating a mechanical signal well beyond the initial activated region via tension and rest-length dynamics (Fig. 2a). The permanent, history-dependent changes accumulated across the entire network result in a stable, reprogrammed mechanical state, which we quantify by measuring the linear shear modulus (*G*) of the fully relaxed tissue. This is done by applying an infinitesimal simple shear and evaluating the second derivative of the energy at zero strain (see SI text for details).

We report the change in shear modulus, Δ*G* = *G*_final_ −*G*_initial_, after a fixed number of active contractile pulses. The direction of the global change (Δ*G*) is determined by the imbalance of the local remodeling rates, 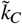 and 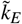, as shown in the phase diagram in Fig. 2b. This ratio serves as the global control knob for programming the tissue’s stiffness. Specifically, for 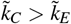, contraction-side remodeling dominates, and the tissue exhibits stiffening (Δ*G >* 0). Conversely, for 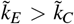, extension-side remodeling dominates, leading to tissue softening (Δ*G <* 0). In the passive limit where remodeling is absent 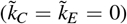, Δ*G* = 0, confirming that the observed changes arise solely from tension remodeling. Since real epithelial tissues are observed to be in the regime 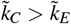[33, 34], *we predict a strain-stiffening behavior under localized active pulses. This offers a concrete, testable hypothesis using optogenetic tools combined with rheological experiments*.

*The magnitude of* Δ*G* is highly tunable. Under a fixed number of pulses, |Δ*G*| increases approximately linearly with the size of the activated region for small *R/L* (Fig. 2c). At fixed *R*, |Δ*G*| also grows with pulse amplitude Γ_*a*_(Fig. 2d). Thus, localized contractions provide a control knob to program global mechanical properties, stiffening or softening, by tuning the balance of remodeling rates and the spatial extent and strength of the driven region.

### Long-range mechanical memory drives cooperative response

Having established that local tension remodeling enables mechanical memory encoding and can reprogram global stiffness, a key question is how this memory is communicated across the tissue, especially over distances exceeding cell size.

To address this, we designed a two-stage simulation. We first applied a localized contraction pulse to a “trained” region (Region 1) of the tissue and allowed the system to mechanically relax. We then applied a second, identical pulse to a distant, “test” region (Region 2), separated by distance *D* (Fig. 3a, Movie S2). We quantify the memory effect by comparing the average active tension generated in Region 2 under two conditions: (1) when Region 1 was previously activated (green curve, Fig. 3b) versus (2) when Region 1 was naive (black curve, Fig. 3b). The results show that the average tension in Region 2 rises more rapidly and reaches a higher final value when the tissue has previously experienced an active contraction elsewhere (Fig. 3b). We define the cooperative enhancement (ΔΛ) as the difference in the final average active tension (Λ) in Region 2 due to the prior activation of Region 1 (Fig. 3b). This non-local change indicates a longrange mechanical memory effect, not previously appreciated in eptihelial mechanics.

**FIG. 3.**
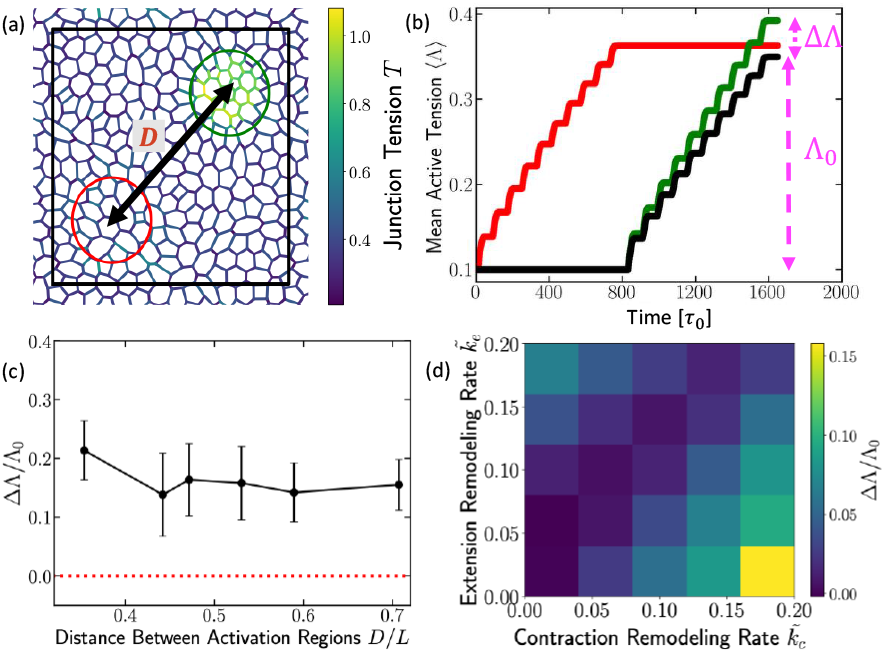
Long-range cooperative effects from local contractile signals. (a) Simulation snapshot showing contractile activations applied to two separate circular regions of the tissue: the first region (red) and the second region (green), each with radius R/L = 1/6. The applied signal matches that of Fig. 1, with 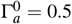, Γ_a_ = 2, and pulse period 40. (b) Average active tension in the first region (red) and in the second region (green) during activation. The black curve shows the average active tension in the second region when the first region was not previously activated. (c) Change in the average active-tension in the second region, computed as the difference between the final values of tensions of the green and the black curves in (b). We refer to this as the cooperative enhancement which is plotted as a function of the center-to-center distance between these two active regions. (d) Cooperative enhancement phase diagram as a function of 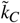 and 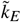.

To determine the spatial extent of this memory, we systematically varied the center-to-center distance *D* between the two activated regions (Fig. 3a). As shown in Figure 3c, the memory effect (ΔΛ) decays slowly with increasing distance *D* but critically saturates to a finite, non-zero value. This suggests that the interaction is not purely local, such as through direct cell-cell contact, but is instead mediated through long-range mechanical coupling in the whole tissue. The permanent tension and rest-length changes stored locally in Region 1 globally bias the elastic response of the tissue, making the distant Region 2 more susceptible to remodeling upon activation.

The strength of this cooperative effect further depends on the rates of contraction 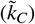and extension 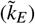 induced tension remodeling (Fig. 3d). This dependence shows that the 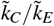 ratio not only dictates the direction of adaptation but also amplifies the magnitude of the long-range memory effect. Specifically, for the experimentally relevant regime where 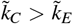, increasing 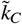 at a fixed 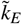increases the coopera-tive enhancement (ΔΛ) (Fig. 3d). This amplification occurs because the initial stimulus (Region 1 activation) creates a large number of high-tension, memory-encoding junctions. A higher 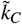ensures that the local tension accumulated in these junctions is greater and more stable, resulting in a globally stiffer initial state for the tissue. When Region 2 is then activated, the enhanced stiffness transmits a stronger baseline stress through the network, pushing the local strain in Region 2 more quickly past the remodeling threshold and accelerating its own tension accumulation.

### Programming auxeticity via cyclic deformations

Having established that tissues can adapt under local contractile activity, we next ask whether such adaptive dynamics can be harnessed to design materials with targeted macroscopic properties. A particularly useful property to train is the Poisson’s ratio (*ν*), which quantifies the transverse response of a material under uniaxial strain. Materials with a negative Poisson’s ratio (auxetic materials) expand laterally when stretched and contract laterally when compressed. These desirable auxetic responses, which enhance shear resistance and energy absorption, are actively pursued in the design of mechanical metamaterials and architected solids [12, 52–54]. Recent studies have shown that periodic mechanical training, cyclically deforming disordered networks, can tune their internal structure to produce auxetic behavior through local remodeling [12]. Here we show that epithelial-like adaptive networks can achieve the same effect autonomously, through mechanosensitive tension remodeling at cell junctions.

To test this programmability, we subject the tissue to externally imposed oscillatory bulk deformations of amplitude *ε*_*B*_ (Fig. 4a, Movie S3). During each cycle of isotropic compression and expansion, all cell edges experience local strain, while the adaptive variables, edge tensions Λ_*ij*_ and rest lengths *L*_*ij*_, evolve according to (2) and (4). These deformations act as a mechanical training protocol that gradually tunes the microscopic edge properties and, consequently, the emergent elastic response of the tissue. After applying a fixed number of bulk-training cycles, we relax the system to mechanical equilibrium and measure its effective Poisson’s ratio. To compute *ν*, we impose a small uniaxial strain *ε*_*y*_ along the *y*-axis, allow the tissue to relax, and measure the induced transverse strain *ε*_*x*_ along the *x*-axis. The Poisson’s ratio is then calculated as *ν* = − *ε*_*x*_*/ε*_*y*_ (Fig. 4a).

**FIG. 4.**
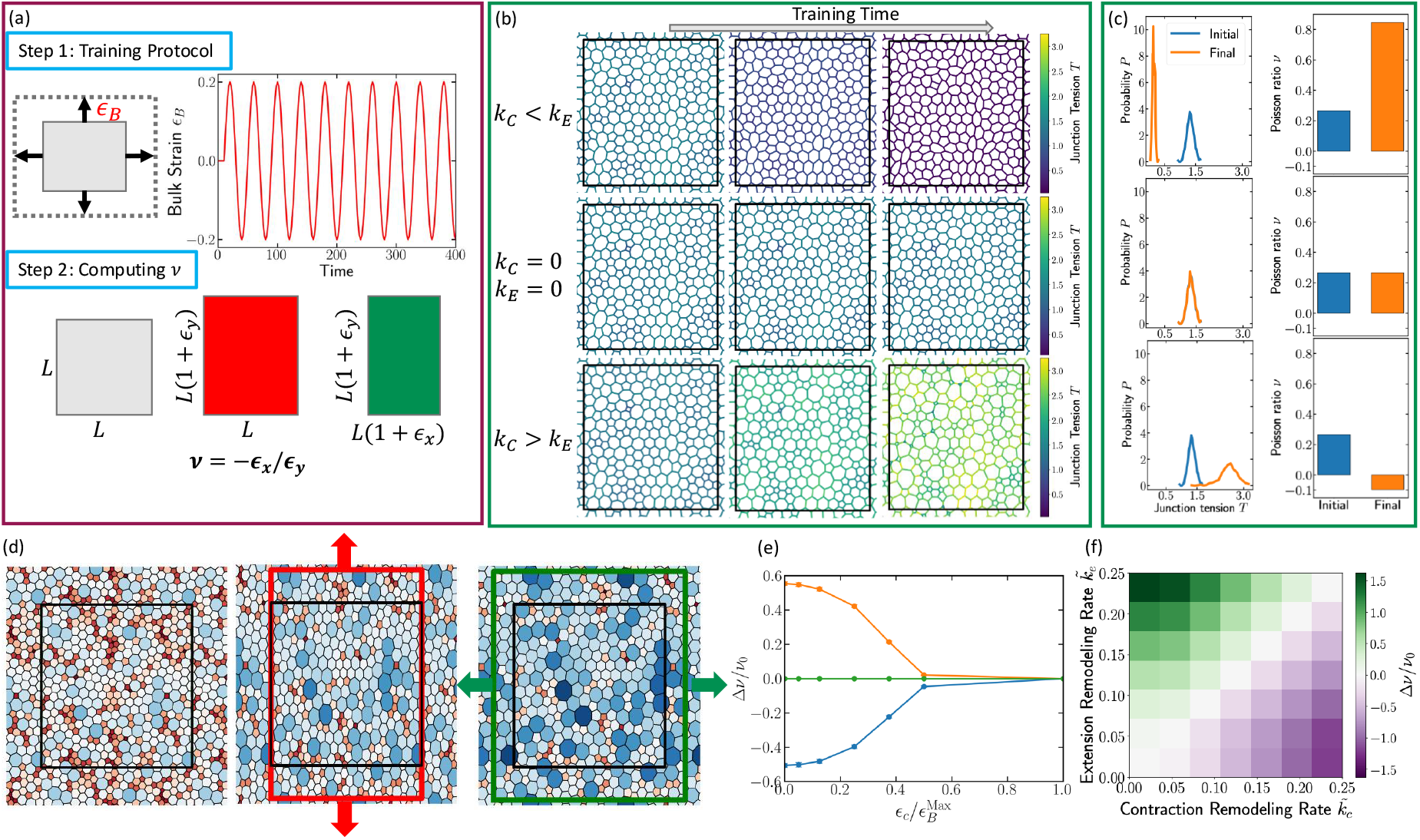
Training auxetic states under periodic bulk deformations. (a) Training tissue’s Poisson ratio under sinusoidal bulk deformations. Upper left panel: schematic of a bulk deformation with magnitude ε_B_. Upper right: sinusoidal bulk-deformation signal applied over time. Lower panel: Schematic of the method used to compute the Poisson ratio. (b) Tissue snapshots under bulk deformation. Top row: evolution from the initial state to mid-training and the final configuration for 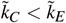. Middle row: case with 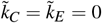. Bottom row: case with 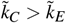. (c) Initial (blue) and final (red) edge-tension distributions for the three cases shown in panel (b). The right three plots show the corresponding initial (blue) and final (red) Poisson ratios for each scenario. (d) Auxetic response of a tissue under uniaxial stretch. Cells are colored by their area deviation, ΔA_α_ = A_α_ − A_0_, with blue indicating larger-than-rest areas and red indicating smaller-than-rest areas. Left: undeformed reference state (black rectangle). Middle: tissue under an imposed deformation along the y-direction (red arrows and red rectange). Right: relaxed configuration showing lateral expansion of the x-boundary (green arrows and green rectangle), consistent with a negative Poisson ratio. For visual clarity, this example uses a relatively large applied deformation; all reported values of the linear Poisson ratio ν are computed using a small strain of 1%. (e) Change in Poisson ratio as a function of the ratio of the critical edge-level strain to the maximum applied bulk strain, 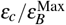. Orange: 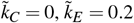; green: 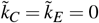; blue: 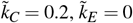. (f) Phase diagram for the case ε_c_ = 0.01 and 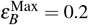.

Figure 4b shows representative tissue configurations and tension fields for three regimes: (1) *k*_*C*_ *< k*_*E*_, (2) *k*_*C*_ = *k*_*E*_ = 0, and (3) *k*_*C*_ *> k*_*E*_. When *k*_*C*_ *< k*_*E*_, edge tensions decrease over repeated bulk cycles, leading to more uniform cell areas (see SI for area distributions) and an increase in *ν*. In the passive control model without remodeling (*k*_*C*_ = *k*_*E*_ = 0), both tensions and *ν* remain unchanged. By contrast, when *k*_*C*_ *> k*_*E*_, edge tensions grow monotonically during training, generating strong structural and tension heterogeneity with coexisting large and small cells (Fig. 4b,c). In the SI, Fig. S3 shows that the area distribution in this regime broadens substantially, with distinct populations of very small and very large cells. Fig. S4 of SI shows that cell pressure distributions narrow and shift to low pressures for *k*_*C*_ *< k*_*E*_ but remain broad, with both negative and positive pressures, for *k*_*C*_ *> k*_*E*_. This highly tensed, heterogeneous architecture drives *ν* downward, ultimately yielding negative values characteristic of an auxetic response (Fig. 4c).

The physical origin of this auxetic behavior is a rigidity switch controlled by the competition between junctional tension and cell area elasticity. In the extension-dominated regime 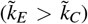, junction tensions are low, and the stiff area elasticity term (*K*_*A*_(*A*_*α*_− *A*_0_)^2^) dictates the mechanical response. To maintain cell area when stretched along the *y*axis, the tissue contracts laterally, resulting in a positive Poisson ratio (*ν >* 0). Conversely, in the regime 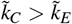, edge tensions increase continuously, and the tension energy term becomes dominant. In this highly tensed state, the cell area deformations becomes effectively softer than the tension network. When subjected to uniaxial stretch (*ε*_*y*_ *>* 0), the vertices surrounding the stretched cells move outward in the transverse direction, causing lateral expansion (*ε*_*x*_ *>* 0) to relieve the high internal tension. This outward motion yields the required negative Poisson ratio (*ν <* 0) (see Fig. 4d).

The 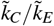 ratio acts as the primary learning bias, as shown in the phase diagram (Fig. 4f), which demonstrates that *ν* can be tuned upward or downward by varying this ratio. Additionally, the training is gated by the ratio of the applied strain to the local remodeling threshold (*ε*_*B*_*/ε*_*c*_). When the applied deformation *ε*_*B*_ is too small, the tissue is inert; as the ratio *ε*_*B*_*/ε*_*c*_ increases, adaptive remodeling becomes stronger, and the magnitude of the programmed change in *ν* grows rapidly (Fig. 4e).

These results demonstrate that epithelia-like adaptive networks can be trained through repeated bulk deformations to acquire programmable elastic responses, including negative Poisson’s ratios, without any designed geometry or plastic flow, purely through dynamic mechanosensitive feedback. In the regime *k*_*C*_ *> k*_*E*_, our theory suggests that under oscillatory external deformations, tissues would continuously stiffen and decrease their Poisson ratio. This presents an intriguing experimentally testable prediction for epithelial monolayers.

## DISCUSSION

A minimal, mechanosensitive tension–remodeling rule is sufficient to produce three hallmark features of trained materials in epithelial sheets: history-dependent adaptation of global moduli, long-range mechanical memory, and learned elastic responses under cyclic driving. The same local update rule that captures optogenetic single-junction behavior [28, 33] reprograms tissue-scale elasticity. In particular, repeated bulk deformations tune the Poisson ratio, including into the auxetic regime, without architected geometry or plastic hinges, demonstrating a functional parallel with the programmability and history dependence observed in directed-aging and periodic-training strategies for disordered solids [12, 55]. This places epithelial mechanics within a broader physical-learning framework [20] in which fast elastic fields drive slow, local parameter updates that accumulate history and alter subsequent response.

The model identifies two primary control levers for programming the material. First, the imbalance between contraction- and extension-side remodeling sets the learning bias and the direction of training. When contraction-driven remodeling rate is higher (*k*_*C*_ *> k*_*E*_), localized pulses stiffen the material and cyclic bulk driving lowers *ν*, switching the material to auxetic behavior. When extension-induced remodeling rate is higher (*k*_*E*_ *> k*_*C*_), the opposite trends occur. Second, the amplitude of the applied strain relative to a local strain threshold for junctions (*ε*_*B*_*/ε*_*c*_) gates learnability: for *ε*_*B*_ *< ε*_*c*_, the epithelial sheet is inert; above threshold, per-cycle updates add coherently and the trained response grows.

Our main results are based on the limit of negligible tension relaxation 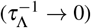, assuming stable, persistent memory storage. To confirm that the observed phenomena are indeed contingent upon this memory, we analyzed the effect of finite tension relaxation timescale on the tissue’s adaptive response (SI Fig. S6). We find that memory-based effects, including the change in shear modulus (Δ*G*), the cooperative long-range response (ΔΛ), and the change in Poisson ratio (Δ*ν*), are all diminished when tension relaxation is fast compared to junction remodeling rate 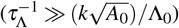, where *k* denotes the strain-dependent remodeling rate (*k* = *k*_*C*_ for contractile strains and *k* = *k*_*E*_ for extensional strains). Only when tension relaxation is slow does the tissue retain a significant, persistent memory and exhibit programmed mechanical changes. This suggests that the remodeling rule constitutes a functional storage mechanism, as the rapid erasure of Λ_*ij*_ eliminates all history-dependent mechanical responses.

Our work proposes new possible experimental tests on epithelial tissues. The tissue global rheology should be programmable by varying the radius and strength of the activated region: the change in shear modulus is predicted to scale approximately linearly with the size of the activation zone for small *R/L* and to grow with pulse amplitude, with the sign determined by *k*_*C*_*/k*_*E*_. The observed long-range mechanical memory also suggests a concrete experimental test: a local input pulse should bias the response to a later, distant pulse after full relaxation, revealing history-dependent coupling stored in cell junctional states. The magnitude of this bias should decay with separation before saturating, reflecting elastic coupling across the sheet. Moreover, under small-amplitude isotropic stretch–compression cycles, the Poisson ratio should change monotonically over training time and reach negative values when *k*_*C*_ *> k*_*E*_.

These behaviors can be probed using optogenetic myosin activation tests and Poisson’s ratio measurements under cyclic training [24, 28, 56, 57]. The predicted tissue stiffening and shift toward auxetic behavior under cycling aligns with concurrent experimental work showing that cyclic loading increases epithelial tissue lifetime and tolerance to large deformations [57]. Measuring the change in Poisson ratio before and after such cyclic loading in epithelial sheets would provide a direct, comprehensive experimental test of our adaptive remodeling mechanism.

## ACKNOWLEDGMENTS

This research was supported in part by the National Institutes of Health (NIH R35 GM143042).

## SUPPLEMENTARY INFORMATION

### Initial tissue configuration

To generate the initial tissue geometry, we use the open-source `CellGPU` package [44]. We place *N* random points as cell centers in a square simulation box of lateral size 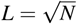 with periodic boundaries, and construct the corresponding cellular network using a Voronoi tessellation. This yields a disordered confluent tissue composed of polygons with various numbers of neighbors.

To obtain a mechanically relaxed reference state, we minimize the classical vertex-model energy

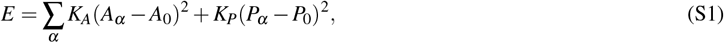

with *K*_*A*_ = *K*_*P*_ = 1, *A*_0_ = 1, and *P*_0_ = 3.7, placing the system in the solid-like regime. Here *A*_*α*_ and *P*_*α*_ denote the instantaneous area and perimeter of cell *α*.

Each cell *α* is represented as an *n*_*α*_-sided polygon with ordered vertices 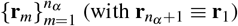. Its geometric measuresare computed as

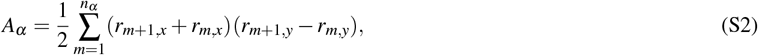

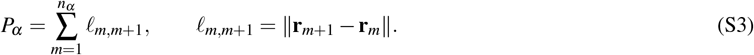

We then relax the tissue using the FIRE algorithm [58] to obtain the minimum-energy configuration. T1 transitions are allowed during this process and are triggered whenever an edge contracts below a threshold length ℓ_*T*1_ = 0.05.

### Shear modulus calculation

To compute the shear modulus *G* of the dynamic tension–remodeling model, we first arrest all dynamical updates of the edge tensions Λ_*ij*_. For a given tissue configuration, the system is then relaxed to a mechanically stable state by minimizing the tension–remodeling vertex-model energy

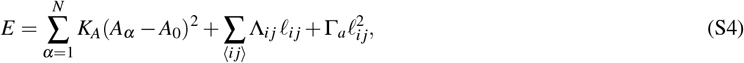

where *A*_*α*_ is the area of cell *α*, ℓ_*ij*_ is the length of edge *i j*, and Λ_*ij*_ are the fixed (frozen) edge tensions at the moment dynamics are halted. The minimization is performed using the FIRE algorithm, following the same above procedure used for preparing relaxed initial states.

Once the mechanically stable configuration is obtained, we compute the shear modulus by evaluating the second total derivative of the energy with respect to an infinitesimal simple shear strain *γ*. For each configuration, we apply an affine simple shear of the simulation box with amplitudes *γ* = − Δ*γ*, 0, +Δ*γ*, updating all vertex positions accordingly. After each shear step, the vertex positions are relaxed using the FIRE algorithm while keeping Λ_*ij*_ fixed, and we record the corresponding energies *E*(− Δ*γ*), *E*(0), and *E*(+Δ*γ*). The shear modulus is then obtained from a centered finite-difference approximation to the second derivative,

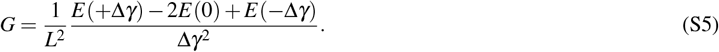

We have verified that the resulting *G* is insensitive to the choice of a small Δ*γ*, confirming that the calculation is performed in the linear elastic regime. We note that this numerical procedure is consistent with the standard definition of the linear shear modulus obtained from the Hessian of the system [59, 60]. Because the tension degrees of freedom are held fixed during these minimizations, this procedure isolates the instantaneous elastic response of a given trained state of the tissue.

### Extension of an edge under contractile signals

In the main text, we showed that when 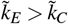, an edge subjected to pulsatile contractile signals can surprisingly undergo a net extension over time. Figure S1 illustrates this behavior. During each contractile pulse, the edge initially shortens, as expected.

**FIG. S1.**
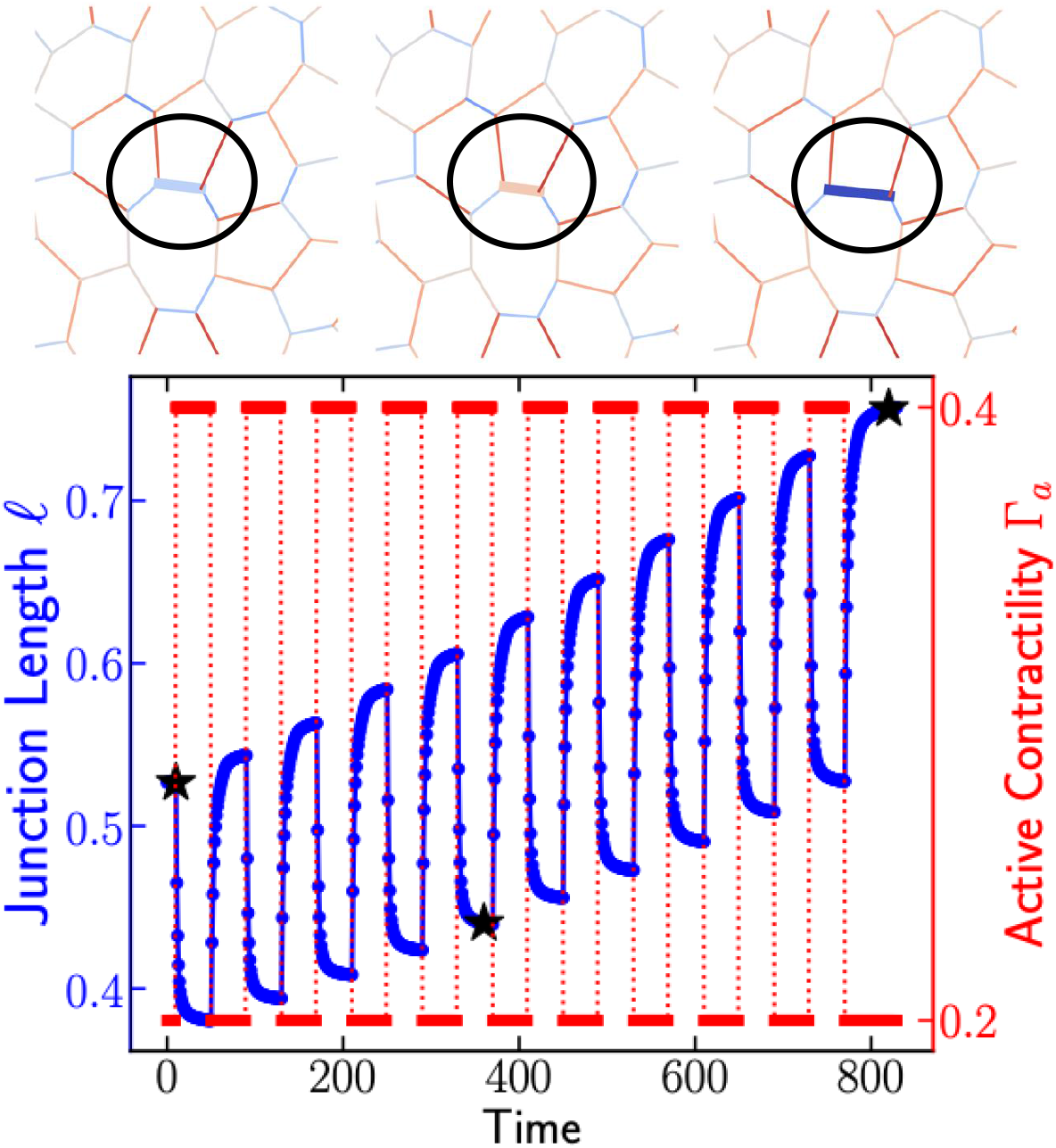
Edge length (left, blue axis) and applied contractile signal (right, red axis) as functions of time in the regime 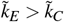 (here 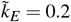 and 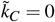). The top row shows consecutive snapshots of the edge corresponding to the time points marked with black stars in the main plot. Here, the junction-level critical strain is *ε*_*C*_ = 0.1.

However, in the relaxation phase that follows, the influence of neighboring edges places the junction in a strain regime where *ε > ε*_*C*_. In this regime, the tension–remodeling rule reduces the edge tension. Repeated cycles of contraction and relaxation therefore drive a progressive decrease in tension, leading to a net extension of the edge despite the applied contractile signals.

### Effect of active contractility period

The duration of the active contractile pulse strongly influences the edge response. Figure S2b shows the net change in edge length as a function of the contractility period *T*. For very short periods, the edge has insufficient time to remodel, resulting in negligible length change. As *T* increases, the accumulated remodeling leads to a monotonic increase in length change, which eventually saturates to a plateau. This has been observed in experiments of epithelial junctions cite.

**FIG. S2.**
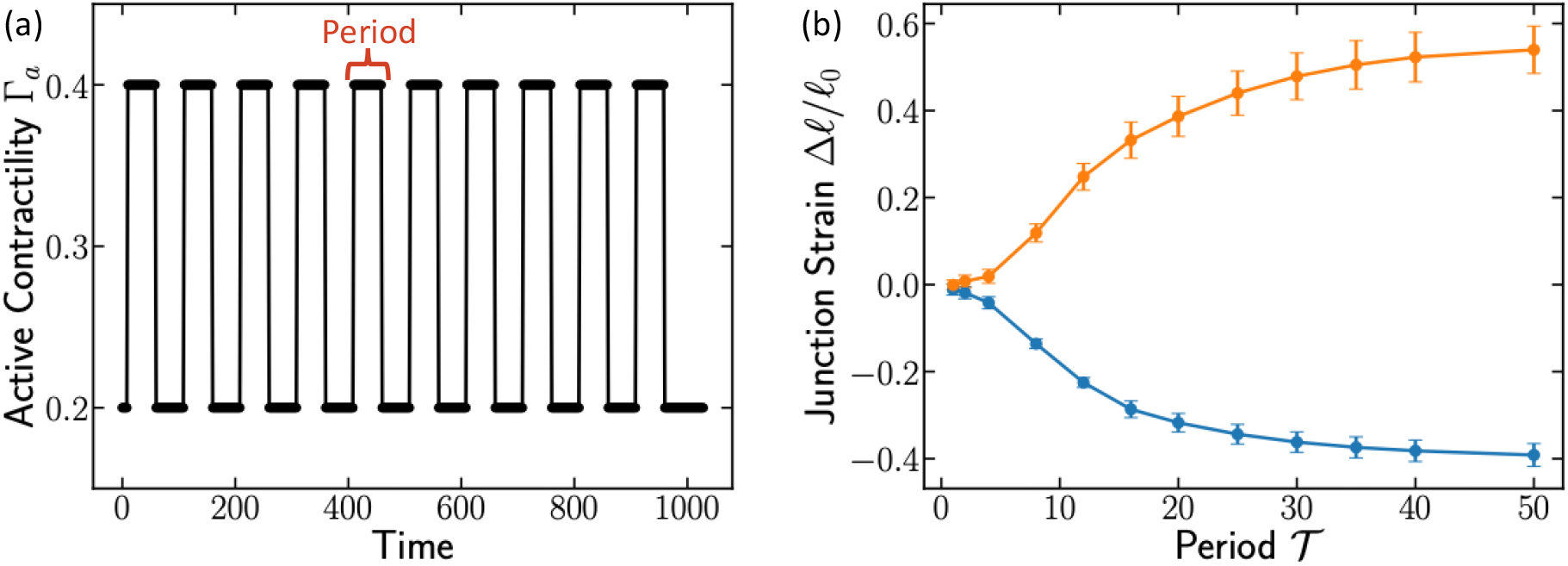
(a) Pulsatile contractility signal alternating between a low value Γ_*a*_ = 0.2 and a high value Γ_*a*_ = 0.4. The period of this signal controls the extent of junctional remodeling. (b) Final change in junction length after applying the signal in (a) for different periods *T*. The orange curve corresponds to the regime 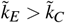 (here 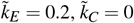), where edges exhibit net extension. The blue curve shows the regime 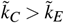 (here 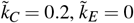), where edges contract under repeated pulses.

### Structural features

Periodic bulk oscillations modify tissue structure through the adaptive coupling between junctional strain and tension. We quantify these structural changes by tracking the evolution of cell-scale geometric statistics after training. As shown in Fig. S3a, when 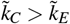 cell areas develops a pronounced high-variance distribution, characterized by the simultaneous presence of cells with enlarged and shrunk areas. This increase in area heterogeneity results in the reduction in Poisson ratio reported in the main text. In contrast, when 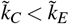, the area distribution narrows and concentrates around the mean area of 1, consistent with a more uniform confluent monolayer.

**FIG. S3.**
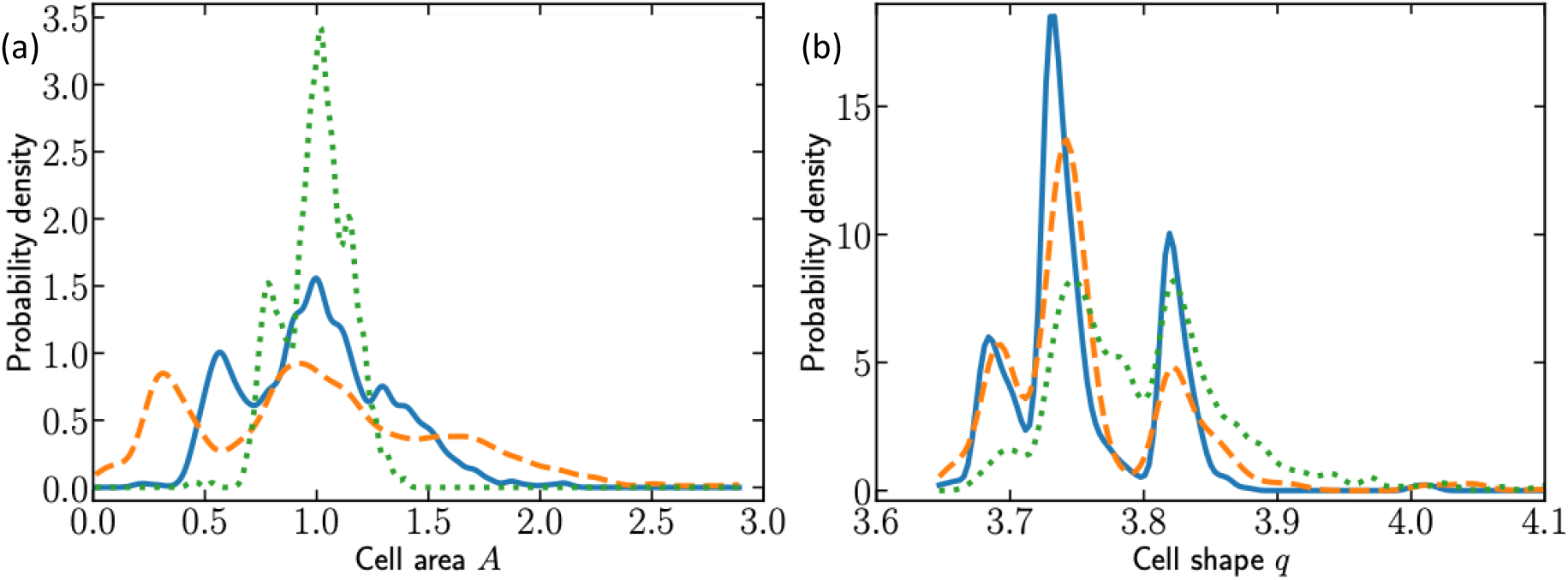
(a) Probability density of cell areas. Shown are the initial distribution (solid blue), the post-training distribution for bulk–strain training with 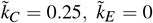 (dashed red), and the post-training distribution for 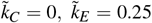 (dotted green). (b) Corresponding distributions of the cell shape index 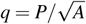 computed from the same configurations as in panel (a). The data is averaged over 5 different random samples.

Figure S4 shows the evolution of cell-level pressure distributions under bulk–strain training. For 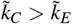, the distribution develops a pronounced both positive high-pressure and negative high-pressure tails, indicating the emergence of tiny and big cell populations as training progresses. In contrast, when 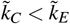, cell pressures shift toward low but positive values, reflecting a more homogeneous cell area distributions. The resulting state is closer to a fluid-like, weakly compressed configuration, consistent with the increase in Poisson ratio reported in the main text.

**FIG. S4.**
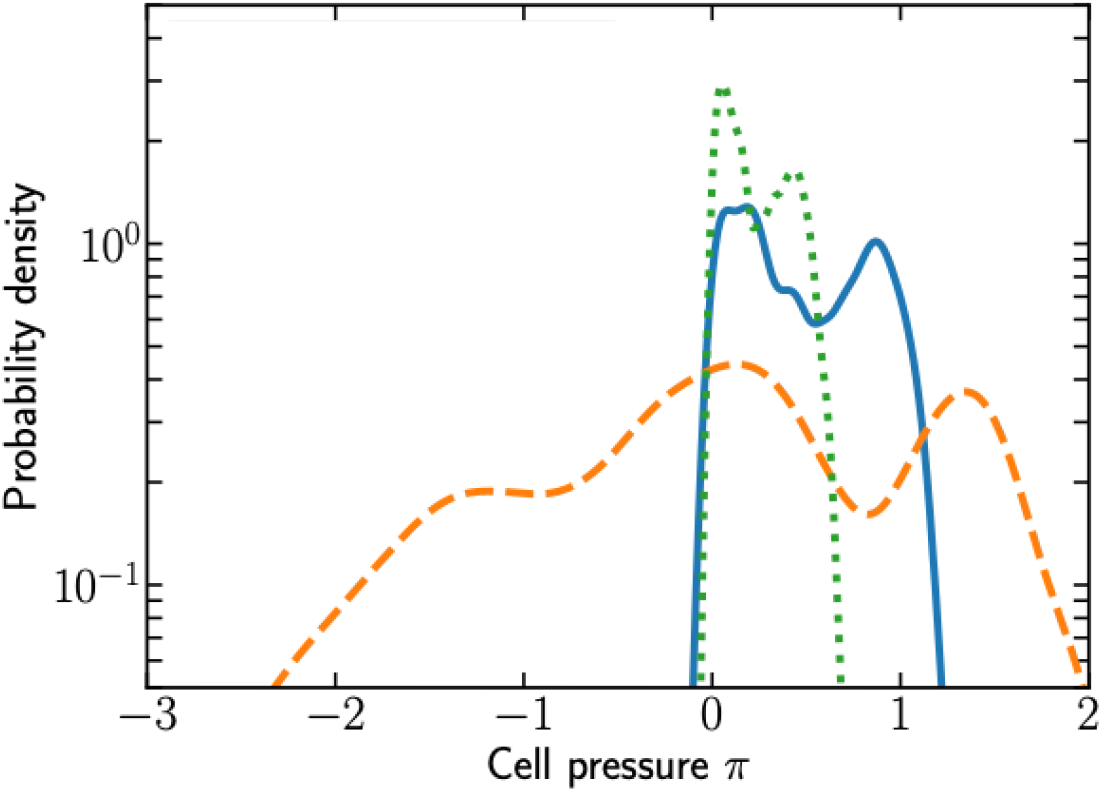
Probability density of cell pressures *π*_*i*_ = − 2*K*_*A*_(*A*_*i*_ − *A*_0_) for the same configurations shown in Fig. S3. Displayed are the initial pressure distribution (solid blue), the post-training distribution for bulk–strain training with 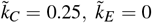 (dashed red), and the post-training distribution for 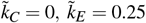 (dotted green). Data are averaged over 5 independent random samples.

Figure S5 shows the distributions of edge strain after bulk–strain training. The initial configuration has zero strain by construction. When 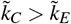, the post-training distribution develops a pronounced heavy tail, indicating the presence of edges that undergo large relative elongation. In contrast, for 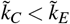, the strain distribution remains sharply peaked, reflecting a population of edges that experience only small deviations from their reference lengths.

**FIG. S5.**
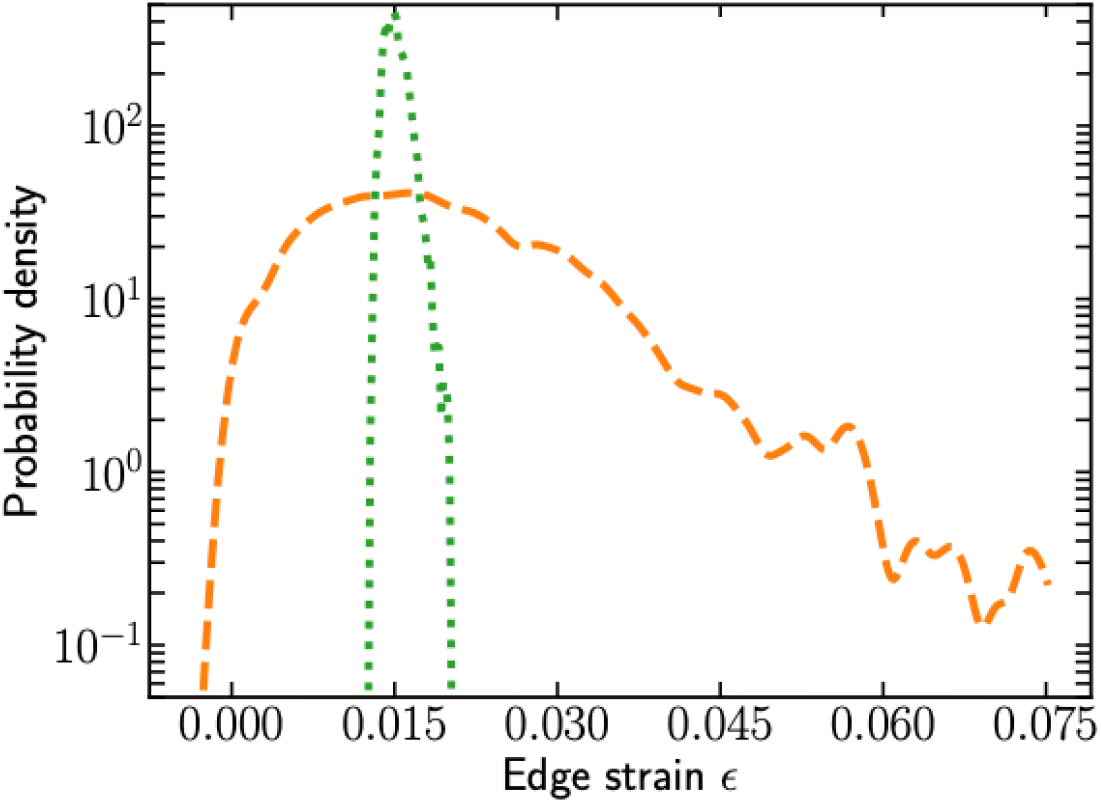
Probability density of edge strains 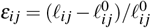 for the trained tissues shown in Fig. S3. Displayed are the post-training strain distributions for bulk–strain training with 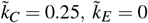 (dashed red) and for 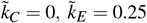 (dotted green). The initial configuration is not shown because all edges begin at zero strain. Data are averaged over 5 independent random samples.

### Effect of finite tension relaxation

The tension–remodeling framework described in the main text generates clear signatures of mechanical memory and adaptive response. In biological tissues, however, junctional tension relaxes over a finite timescale due to actomyosin turnover and adhesion dynamics. Rapid relaxation should diminish the influence of prior loading, whereas slow relaxation preserves it. All results presented so far correspond to the limit of no relaxation, *τ*_Λ_ = ∞.

To examine how finite relaxation alters the dynamics, we augment the tension update rule by adding a relaxation term:

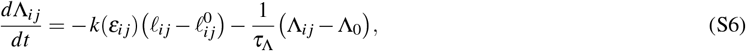

where the first term is the strain-dependent remodeling introduced in the main text, and the second term drives Λ_*ij*_ toward a baseline value Λ_0_ over a timescale *τ*_Λ_.

To examine how tension relaxation shapes mechanical memory, we first measured how a localized contractile pulse alters the shear modulus. Figure S6a shows the resulting change in shear modulus as a function of the inverse relaxation timescale 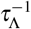. When relaxation is fast, the imposed tension rapidly returns to its baseline value and the tissue retains no memory of the pulse, leading to essentially no change in *G*. As relaxation slows (smaller 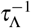), the effects of the contractile event persist, resulting in a finite increase in the shear modulus. In the limit of very slow relaxation, 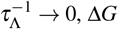 approaches the no-relaxation behavior described in the main text.

We next quantified how finite tension relaxation modifies the cooperative, long-range memory effect described in the main text. In the absence of relaxation, tissues that previously experienced a contractile pulse in one region respond more strongly to a subsequent pulse applied in a distant region, reflecting persistent changes in the underlying tension field. To measure this cooperative effect, we computed the difference between the final average tension in the second activated region when a prior contractile event occurred elsewhere and the corresponding tension when no prior activation was applied. Figure S6b shows the normalized response ΔΛ*/*Λ_0_ in the second region as a function of the inverse relaxation timescale 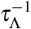. When relaxation is fast (large 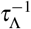), the cooperative effect vanishes: tension rapidly returns to its baseline value, eliminating any imprint of earlier contractile signals. As relaxation slows (smaller 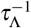), the cooperative response increases, indicating the accumulation of a finite mechanical memory. In the limit of very slow relaxation, the effect saturates to a plateau corresponding to the no-relaxation regime.

Figure S6c shows how the Poisson’s ratio changes in tissues trained under the bulk oscillation protocol described in the main text, plotted as a function of the inverse tension–relaxation timescale 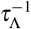. When tension relaxes rapidly larger 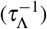, the system effectively erases the history of loading, and the final mechanical state remains unchanged. In contrast, slow relaxation allows tension updates to accumulate, producing a finite shift in tissue properties and a persistent trained state. In biological epithelia, tension relaxation is governed by actin turnover and myosin motor dynamics, which operate on finite timescales; as a result, tissues can maintain mechanical memory over repeated deformation cycles.

**FIG. S6.**
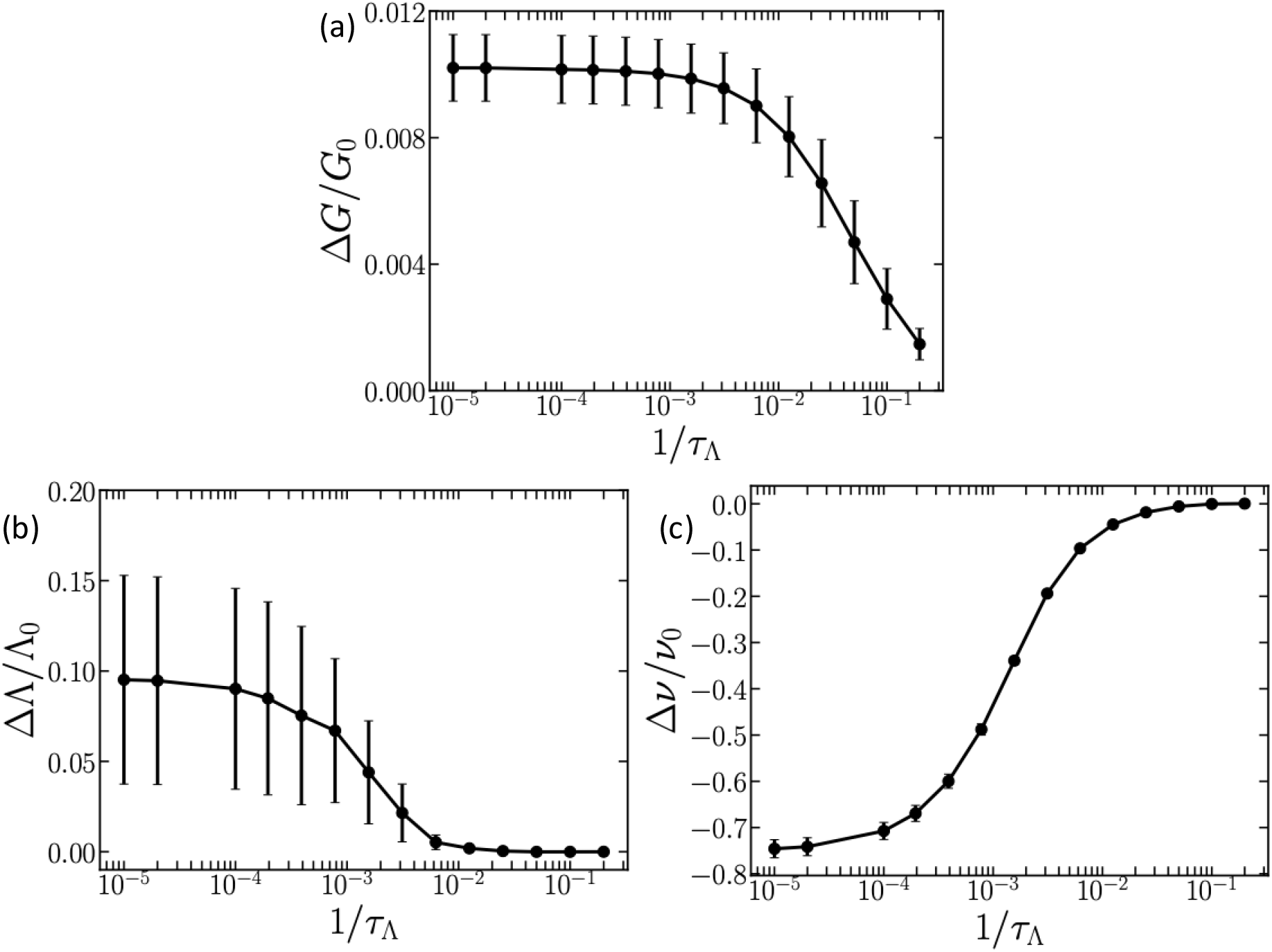
(a) Normalized change in shear modulus Δ*G/G*_0_ as a function of the inverse tension–relaxation timescale 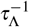. The tissue is trained by applying a single contractile pulse in a central circular region of radius *R* = *L/*6 (with 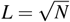 and *N* = 200 cells). The active signal is increased from Γ_*a*_ = 0.5 to Γ_*a*_ = 2.0 for a duration *T* = 40. Results are shown for contraction and extension remodeling rates *k*_*C*_ = 0.2 and *k*_*E*_ = 0.05. (b) Cooperative tension–memory response ΔΛ*/*Λ_0_ in a distant region as a function of 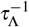. The contractile signal is again pulsed between Γ_*a*_ = 0.5 and Γ_*a*_ = 2.0 for a duration *T* = 40, but here applied repeatedly for 10 on–off cycles. The center–to–center separation between the previously activated region and the probe region is *D/L* = 0.53. The same remodeling parameters *k*_*C*_ = 0.2 and *k*_*E*_ = 0.05 are used. (c) Normalized change in Poisson’s ratio as a function of the inverse tension–relaxation timescale 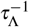, for tissues trained under the bulk oscillatory protocol described in the main text. Results are shown for contraction and extension remodeling rates 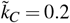 and 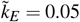. The bulk strain amplitude for this training is *ε*_*B*_ = 0.2 and the edge-level critical strain is *ε*_*C*_ = 0.01.

## SUPPLEMENTARY MOVIES

**Movie S1. Global effects of local contractions**. Time-lapse of a tissue subjected to pulsatile active contractility (Γ_*a*_(*t*)) in a circular region of radius *R* at the tissue center, in the regime *k*_*C*_ *> k*_*E*_. Cell–cell junctions are colored by their total tension *T*_*ij*_. The shear modulus *G*, computed under this activation, increases over time, illustrating the stiffening behavior described in the main text.

**Movie S2. Cooperative effects of local contractions**. Time-lapse of a tissue subjected to pulsatile active contractility (Γ_*a*_(*t*)) in two circular regions of radius *R* separated by a distance *D*, in the regime *k*_*C*_ *> k*_*E*_. Cell–cell junctions are colored by their active tensions Λ_*ij*_. The movie also shows the time evolution of the average active tension in each region, highlighting the enhanced response in the second region due to prior activation of the first, as quantified in the cooperative enhancement analysis in the main text.

**Movie S3. Training tissues under oscillatory bulk deformation**. Time-lapse of a tissue subjected to a sinusoidal bulk deformation signal *ε*_*B*_(*t*) in the regime *k*_*C*_ *> k*_*E*_. Cell–cell junctions are colored by their tension *T*_*ij*_. During training, junctional tensions grow and the tissue develops a heterogeneous structure with coexisting very large and very small cells. The Poisson ratio *ν* decreases over successive cycles and becomes negative, demonstrating the emergence of auxetic behavior discussed in the main text.

